# Projective oblique plane structured illumination microscopy

**DOI:** 10.1101/2023.08.08.552447

**Authors:** Bo-Jui Chang, Douglas Shepherd, Reto Fiolka

## Abstract

Structured illumination microscopy (SIM) can double the spatial resolution of a fluorescence microscope and video rate live cell imaging in a two-dimensional format has been demonstrated. However, rapid implementations of 2D SIM typically only cover a narrow slice of the sample immediately at the coverslip, with most of the cellular volume out of reach. Here we implement oblique plane structured illumination microscopy (OPSIM) in a projection format to rapidly image an entire cell in a 2D SIM framework. As no mechanical scanning of the sample or objective is involved, this technique has the potential for rapid projection imaging with doubled resolution. We characterize the spatial resolution with fluorescent nanospheres, compare projection and 3D imaging using OPSIM and image mitochondria and ER dynamics across an entire cell at up to 2.7 Hz. To our knowledge, this represents the fastest whole cell SIM imaging to date.

## 1. Introduction

Structured illumination microscopy can double the resolution of fluorescence microscopy, is compatible with commonly used fluorophores and sample preparations, requires only modest to medium laser powers and is fast enough to follow cellular dynamics [1, 2]. As such, it has established itself as an important tool in biological fluorescence microscopy. However, the fastest SIM acquisition rates, up to several tens of Hz, have been achieved in 2D SIM, typically in Total Internal Reflection Fluorescence (TIRF) or grazing incidence (GI) modalities to improve optical sectioning [3-6]. As such, imaging is locked to the coverslip-cell interface. While this enables rapid and sensitive imaging of the plasma membrane, it does not allow deeper imaging inside the cell or to visualize the dorsal membrane. 3D SIM breaks free from the coverslip for volume acquisition, at the cost of acquisition speed, due to requiring 15 stacks of different illumination phases and orientations [7, 8]. Processing of a single slice at an arbitrary focal plane has been suggested theoretically [9] and implemented practically [10, 11], and as such can reach faster rates than 3D SIM. However, it only covers a slice limited by the microscope’s depth of focus (<1 micron) and suffers from more background and related artifacts compared to TIRF or GI SIM.

Here we report projection imaging under structured illumination to image an entire cell at acquisition rates faster than 1 Hz. We leverage two recent innovations, oblique plane structured illumination microscopy (OPSIM) [12] and multi-angle projection imaging [13]. Projective OPSIM, or POPSIM for short, enables 2D SIM imaging of projections of an entire cell instead of a thin slice. Because POPSIM projects the volume under structured illumination into a 2D plane, only 9 raw images are required to reconstruct the projected volume. As such, rapid organelle dynamics can be captured throughout a cell, and imaging is not tied to the coverslip-cell interface. Obviously, this comes at the expense of sample information in the third dimension, which is collapsed into a projection. However, for many rapid dynamics, and processes for which simultaneous observation is paramount, projection imaging represents a valuable alternative to slower 3D stacking.

For projection imaging in general, contrast and resolution are often compromised compared to 3D imaging modalities. Thus, structured illumination can provide a mechanism that improves resolving power and/or reduces background haze, depending on the used illumination patterns. While our implementation of projection imaging is specific to a class of light-sheet microscopes that possess rapid scanning mechanisms, some of our results using structured illumination may be generalizable to other projection modalities that are recorded in a widefield format [14-17].

We characterize the imaging performance of POPSIM using fluorescent nanospheres and compare 3D and projection imaging on fixed cells. Finally, we apply POPSIM for imaging the dynamics of mitochondria and the endoplasmic reticulum across entire osteosarcoma cells at up to 2.7Hz acquisition rate.

## 2. Methods

### 2.1 Concept

OPSIM is a combination of light-sheet microscopy and structured illumination, implemented in a single objective format (in the sense that illumination and fluorescence detection occur through the same primary lens) [12, 18]. A schematic representation is shown in Figure 1A. Two mutually coherent light-sheets interfere in a common oblique plane which is inclined by 45 degrees to the optical axis. The light-sheets can be rotated, phase stepped and scanned laterally. In its first conception, three stacks under different phases of the illumination pattern were acquired for three azimuthal orientations of the illumination and detection path [12].

**Fig. 1.**
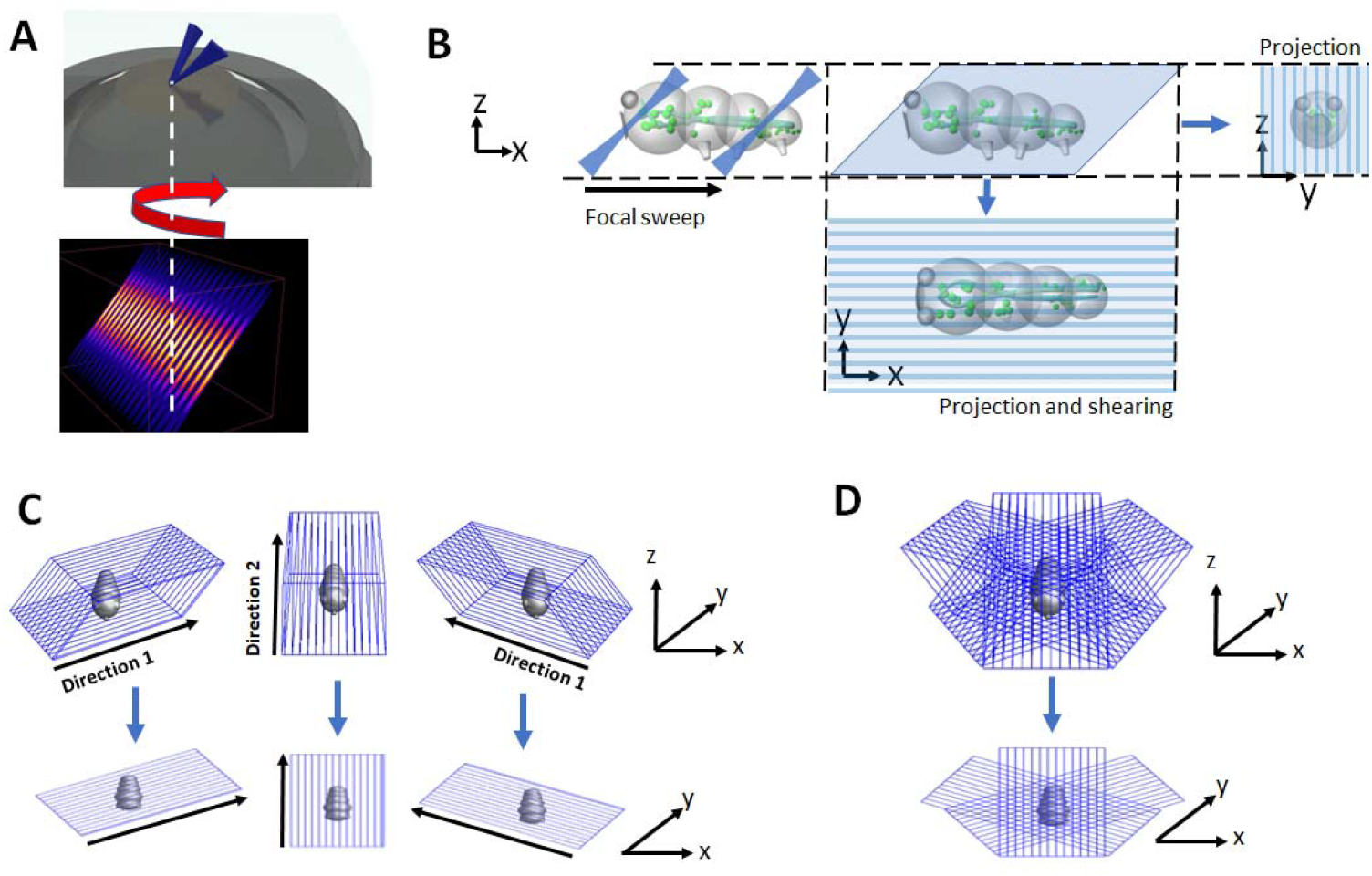
Schematic representation of the POPSIM concept. **A** In OPSIM, a structured light sheet (numerical simulation of the intensity pattern shown below) emerges from a high numerical aperture lens. The structured light sheet can be azimuthally rotated, phase stepped, and scanned laterally to acquire stacks. **B** Schematic illustration of projection imaging. The light-sheet and focal plane are rapidly swept through the sample, covering the sample (light-blue parallelogram). If at least one sweep occurs during a camera exposure, a sum projection is formed (top right). By synchronously shearing the images on the camera during the focal sweep, a projection along the z-axis can be formed (bottom). **C** In OPSIM stacks under different phases of the structured light-sheet are acquired for three azimuthal directions (top). In POPSIM, each volume is projected along the z-axis (bottom). **D** In OPSIM, volumes acquired under different directions are computationally registered in 3D to each other. In POPSIM, the corresponding projections are registered in 2D to each other.

The OPSIM architecture is conceptually compatible with our recently introduced multi-angle projection technique [13]: by rapidly sweeping the structured light-sheet through the sample, and optically shearing the resulting image over the camera detector, a sum projection of the sample is created optically via the shear-warp transform [19], as schematically shown in Figure 1B. We reasoned that if all nine volumes in OPSIM are projected into the same coordinate frame, then SIM processing in a conventional 2D format can be performed. This idea is schematically shown in Figure 1C: volumes that are spanned by rapidly sweeping the structured light-sheet are projected in a “top down” view, i.e., along the normal direction to the coverslip surface (labeled z). The three illumination directions (labeled “Direction 1-3”, also called “SIM orientations” henceforth) are mapped into the lateral x-y plane.

In OPSIM, volumes from the three illumination directions need to be 3D registered to each other. In the projective variant, the registration happens in 2D, as schematically shown in Figure 1D. In contrast to traditional OPSIM, where hundreds of image frames need to be acquired to span a single volume, acquiring only 9 projection images in POPSIM promises a significant reduction in data volume and a corresponding improvement in acquisition speed.

### 2.2 Experimental setup

For the first demonstration of POPSIM, we added a galvo shear unit for projection imaging to our previously published OPSIM instrument. While the details of the OPSIM system have been published elsewhere [12], in short it uses a Nikon NA 1.35 100X silicone oil primary objective and employs structured light-sheets that are tilted by 45 degrees to the optical axis. The structured light-sheets are generated in a Michelson interferometer, and a rapid image rotator is used to switch between three azimuthal orientations of the illumination and detection path. The line spacing of the structured light-sheets can be tuned but faces an upper limit of about 300 nm for an excitation wavelength of 488nm, limited by the numerical aperture of the objective and the off-center position of the illumination beams in its pupil.

In POPSIM, projecting the volumes corresponding to the three SIM orientations into the same 2D plane is crucial. We used a two-step procedure to ensure correct calibration for the sheer galvo unit. First, we performed coarse calibration using a micro-ruler. For a top-down view direction, we expect a √2 stretch in the projection direction given the 45-degree tilt angle of the tertiary imaging system. The shear amplitude was adjusted until the micro-ruler image appeared √2 stretched in the shear direction. Second, we performed fine calibration using two bead-coated coverslips, sandwiched together, with a ∼30-micron gap between the coverslips (See also Figure 2A). We densely coated the bottom coverslip with 100nm fluorescent nanospheres (ThermoFisher, F8803) and the top coverslip sparsely with 500nm fluorescent nanospheres (ThermoFisher, F8813). Subsequently, we filled the gap between the two coverslips with water. While imaging the double-layer coverslip sample, the shear factor was empirically varied in small steps.

**Fig. 2.**
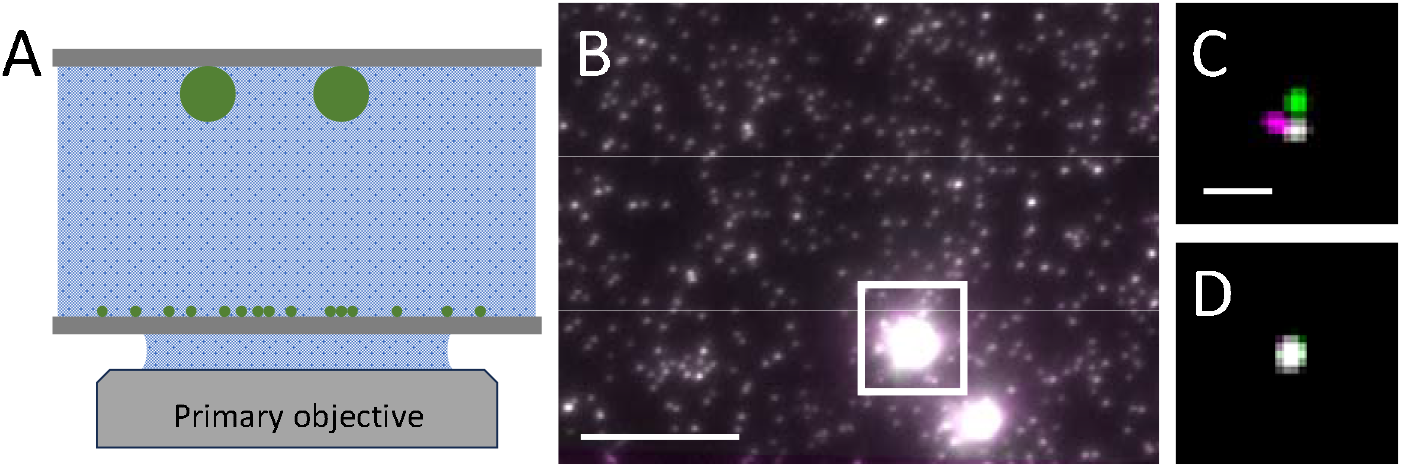
Calibration of the projection viewing angles by mapping two bead layers on top of each other. **A** Schematic illustration of two opposing coverslips, one densely coated with 100nm fluorescent nanospheres on the bottom and one sparsely coated with 500nm nanospheres on top. **B** Overlay of three registered projections corresponding to the SIM orientations 1-3 (color coded green, magenta and gray). The two bright spots are two 500nm beads which are saturated to better show the overlap of the 100nm beads. **C** Inset of bright 500nm nanosphere in the white rectangle in **A**, with contrast adjusted. **D** the same sphere after calibration of the shear parameters. Scale bars: **A**: 10*μ*m; **B**: 2*μ*m.

At each step, we tested if registering the projection images for the three SIM orientations (Figure 1) was successful, i.e., how well the 100nm and 500nm beads overlapped (Figure 2B-D). If the shear amplitude was not optimal, i.e., the resulting projection axis was not the same for each SIM orientation, the registration algorithm could not converge on a solution to simultaneously register the beads projected from the bottom and top coverslip (Figure 2C). The requirement for precise physical registration versus post hoc non-rigid computation registration is a key difference between POPSIM and the original OPSIM implementation.

The process of projection imaging for the double-layer bead sample is instructive in understanding the alignment requirements of POPSIM. If the individual viewing angles differ for the three SIM orientations, the projections physically map the top layer beads onto different locations relative to the bottom layer beads. In the case of mismatched views, computational registration will converge on unique solutions for the bottom or top beads, but not both. For the beads sampled here, our registration algorithm utilized the denser bottom bead layer, leading to divergent features in the top layer if the shearing parameters were incorrect. To complete the fine registration, we iteratively varied the shear parameters in small increments until the bottom and top beads overlapped after registration (Figure 2B & D).

## 3. Results

### 3.1 Resolution measurements

We evaluated the resolution of POPSIM by imaging 100nm fluorescent nanospheres (Polysciences). Figure 3A shows a conventional projection view, which was obtained by summing the nine registered POPSIM frames. This is representative of a projection image that could be obtained by a conventional OPM microscope. Figure 3B shows the same area, but after a SIM reconstruction of the POPSIM data. Chevrons point at pairs of fluorescent nanospheres that were resolved by structured illumination but cannot be readily distinguished in the conventional projection image. We estimated the resolution by measuring the full width half maximum (FWHM) over multiple beads. In Figure 3A, the FWHM was 381±44nm and 386±29nm (mean and standard deviation, n=22) in the x- and y-direction, respectively. This is a lower spatial resolution than what the same microscope can produce in a 3D imaging modality, likely because the 3D PSF is summed along the z-direction.

**Fig. 3.**
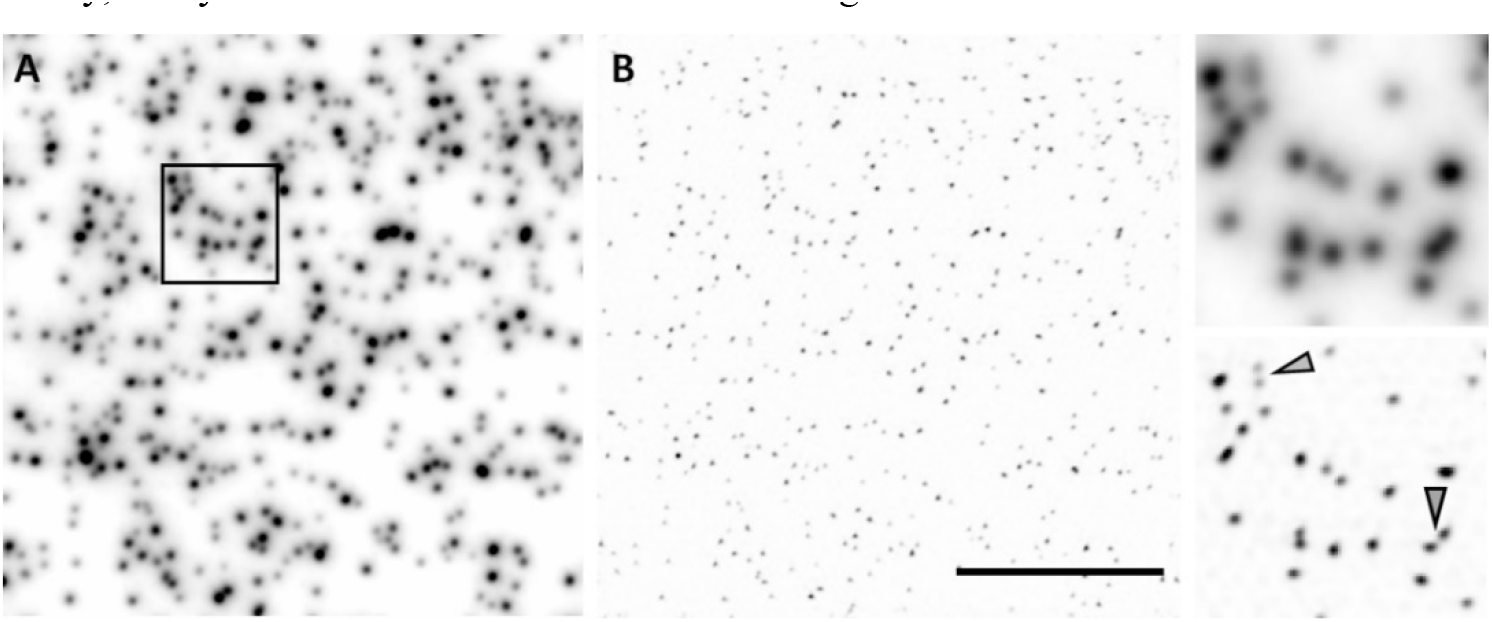
Resolution measurements using fluorescent nanospheres. **A** Projection image of 100nm fluorescent nanospheres obtained by summing all 9 frames of a POPSIM acquisition. **B** POPSIM reconstruction of the same bead dataset. Insets show magnified regions of the black boxed area shown in **A**. Chevrons point at closely spaced beads that are resolved by POPSIM, but not by the conventional projection image. Scale Bar: 10 microns.

The sum projection involves adding portions that are slightly outside of the depth of focus of the imaging system, resulting in slight blurring of the projection PSF. Accordingly, we set the linespacing of the structured light-sheet to ∼390nm, resulting in a frequency that is of similar magnitude as the cut-off frequency of our projection OTF. This linespacing is a bit wider than what practically is possible on our setup (∼300nm at 488nm excitation wavelength). In the SIM mode, we measured an FWHM of 185±10nm and 189±11nm (n=22) for the x- and y-direction respectively using a traditional SIM reconstruction algorithm with Wiener filtering [20]. Replacing Wiener Filtering with 5 iterations of a Richardson Lucy (RL) iterative deconvolution resulted in an FWHM of 161±11nm and 169±9nm (n=22), respectively, while image decorrelation analysis [21] estimated a resolution of 178nm. The RL deconvolution was not used to boost the resolution further. Instead, we found that the RL assisted reconstruction was more robust than the classical SIM algorithm: there was little to no parameter tuning necessary and it also reduces background haze, which can accumulate due to the sum projection in more densely labeled samples. As such, we employed the RL assisted reconstructions to the biological imaging shown below. The code was the same as the RL assisted slice by slice reconstruction used in OPSIM [12].

### 3.2 Cellular imaging on fixed cells

To compare volumetric and projective SIM imaging we imaged a fixed U2OS cell where mitochondria were stained with Alexa 488 using antibody labeling for MIC60. We further leveraged the fact that we can perform OPSIM and POPSIM imaging on the same setup. In Figure 4A, a maximum projection image, color coded for depth, of the OPSIM data is shown. Notably, mitochondria are present at different heights, with two mitochondria in the center being the highest above the coverslip. In Figure 4B, the corresponding POPSIM image is shown. The projection encompasses the entire cell, so mitochondria near the coverslip as well as the mitochondria presumably above the nucleus are captured. In the insets in Figure 4C-D show a comparison between the 3D and projective imaging. Similar morphological detail is captured of a cluster of mitochondria. Generally, there is good correspondence between features and morphologies, with some exceptions (see for example the chevron in Figure 4B where the reconstruction of mitochondria appears blurrier in the POPSIM image).

**Fig. 4.**
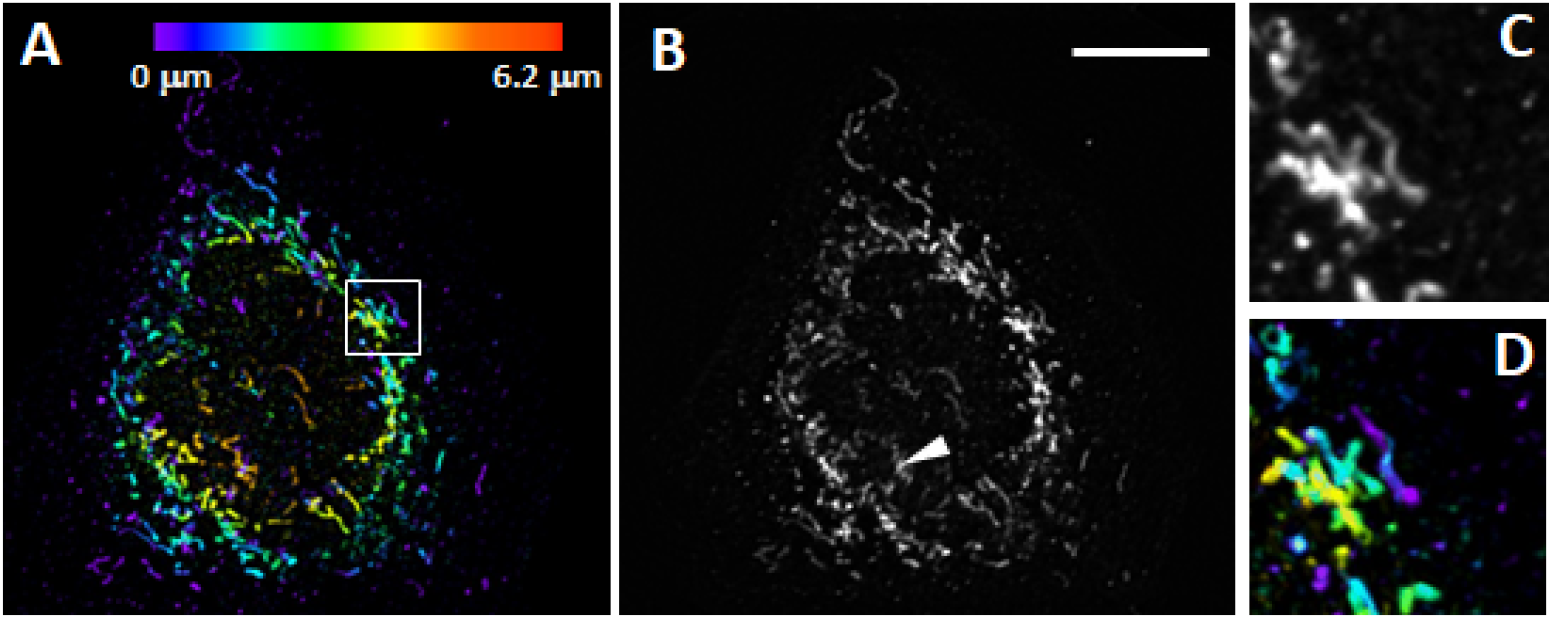
Comparison of 3D and projective imaging. **A** An U2OS cell labeled with MIC60, as imaged with OPSIM. Depth is color coded. **B** The same cell as imaged under POPSIM. The white Chevron points at a reconstruction artifact. **C-D** magnified views of the boxed region in **A**, under POPSIM (**C**) and OPSIM (**D**). Scale bar: 10 microns.

### 3.3 Dynamic live cell imaging

We explored the temporal capabilities of POPSIM by imaging live U2OS cells labeled with OMP-GFP, an outer membrane marker of mitochondria at a rate of 2.2Hz (Figure 5A-C). Subsequent frames showed some jitter artifacts, which may originate from fluctuating edge artifacts the camera had at this speed, or rapid motion of the sample itself. By taking a two-frame average, the artifacts mostly disappeared. The magnified views reveal that POPSIM was able to resolve the hollow structure of the mitochondria (Figure 5C). Since POPSIM performs a sum projection along the z-axis, we expect that the mitochondria walls are less sharply resolved as in a 3D SIM modality, where either a cross-section or a maximum intensity projection can be shown. In Supplementary Movie 1, mitochondria dynamics over 100 timepoints are shown. In multiple instances, rapid protrusion and retraction of mitochondria could be observed.

**Fig. 5.**
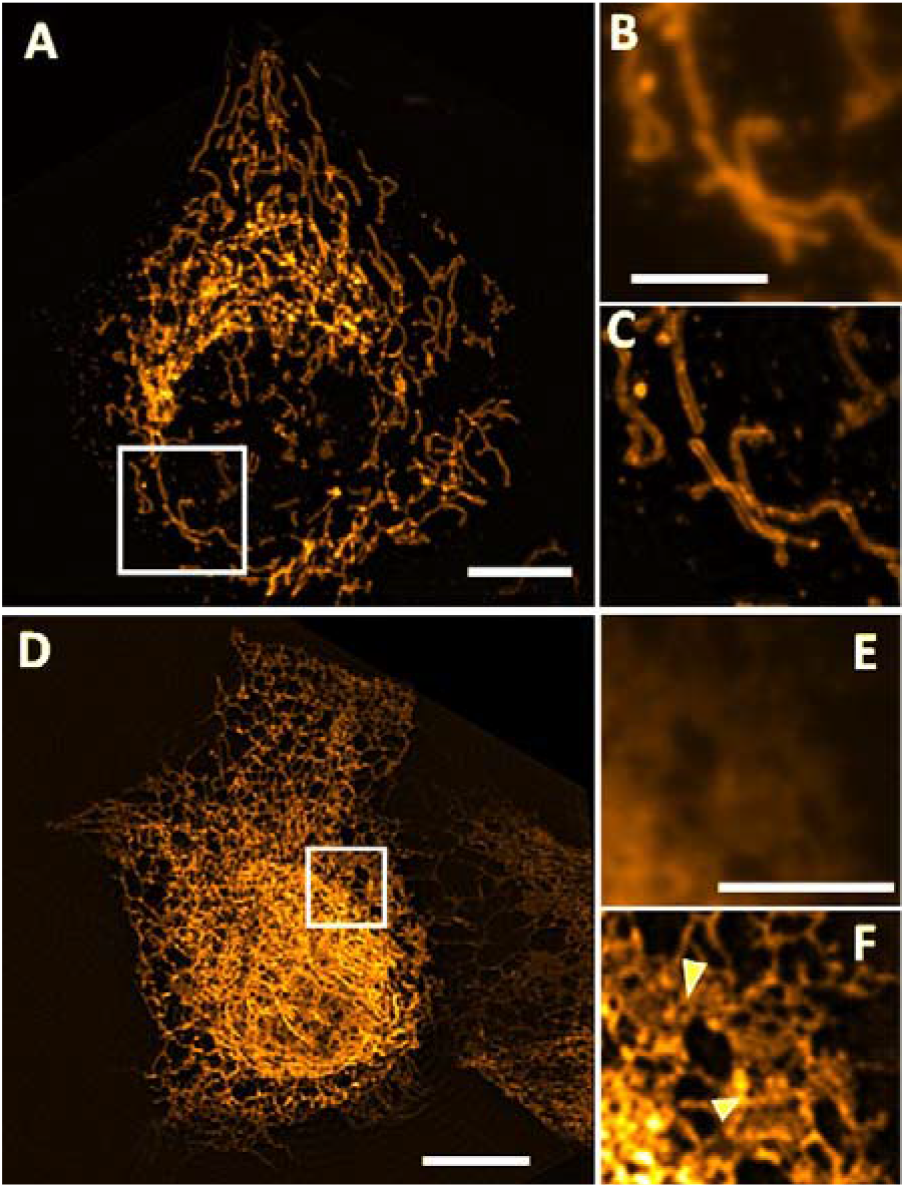
Live imaging with POPSIM. A U2OS cells labeled with OMP-GFP, as imaged with POPSIM at 2.2Hz overall rate. A two-frame average is shown. B Magnified view of the boxed region in A as imaged with projection OPM. C Magnified view of the boxed region in A as imaged with POPSIM. D An U2OS cells expressing the endoplasmic reticulum label Sec61-GFP, as imaged by POPSIM at 2.7Hz overall rate. A two-frame average is shown. E Magnified view of the boxed region in D, as imaged with projection OPM. F Magnified view of the boxed region in D as imaged with POPSIM. Chevrons point to small holes in ER sheet. Scale bars: A,D: 10 microns B,E: 5 microns.

We next imaged live U2OS cells expressing Sec61-GFP, a label for the endoplasmic reticulum (ER) at a rate of 2.7 Hz (Figure 5D-F). POPSIM was able to follow rapid rearrangements of the ER (Supplementary Movie 2). We were also able to resolve fine structures in ER sheets that may represent nanoholes that have been recently described with rapid 2D super-resolution methods [22, 23], albeit the spatiotemporal resolution in our system is lower. These holes persisted in some instances over multiple frames and appeared to move along the sheet. Of note in Figure 5D, the top right corner appears darker than the rest of the image. This edge marks the end of scan direction 3, and the image registration algorithm filled the corner with zeros.

## 4. Conclusions

In summary, we have introduced a rapid way to image entire cells in a projective format with structured illumination. This extends the capabilities of structured illumination to rapidly capture dynamics across an entire cell at rates that cannot currently be achieved with 3D super-resolution methods. Compared to other rapid 2D SIM methods which can only capture a thin slice of 100-1000nm thickness, POPSIM can capture organelle dynamics in a projection of an entire cell. As our imaging was performed on cells with about 10-micron height, a ten-to hundredfold larger volume is sampled than in TIRF- or GI-SIM. This is in our view an important distinction and represents complementary capabilities to other SIM and super-resolution modalities.

Our POPSIM instrument is based on an oblique plane microscope equipped with an NA 1.35 silicone oil objective. As such, it is limited to single cell imaging, due to the limited range over which remote focusing, a key component of OPM, is diffraction limited. Nevertheless, we believe that POPSIM could also be implemented with NA 1.1-1.2 water immersion objectives which possess longer working distances. A separate constraint is how well the shear warp projections map the three SIM orientations into the same plane. This depends on how well the shear unit is calibrated, and how well OPM performs telecentric scanning of a volume.

Simpler implementations of POPSIM with only one SIM direction could be envisioned. This would dispense with the shear calibration and image registration, and light-sheet systems with one illumination direction could be used [24, 25]. Further, it would enable arbitrary viewing directions, compared to the fixed “top down” view employed here, at the expense of anisotropic resolution. Coarser patterns could be applied in this context to increase the optical sectioning strength, which could be valuable in the presence of large background levels.

The Resolution of POPSIM is lower than what SIM systems using sinusoidal interference patterns can achieve. As an example, TIRF-SIM can achieve resolution levels slightly below 100nm. This is in part due to the twice as fine line-spacings of the interference patterns (∼185nm compared to 390nm used here, assuming 488nm excitation wavelength) that can be generated in the evanescent fields, and the smaller detection PSF. Finer line spacings could be generated in POPSIM using higher NA TIRF objectives. Such oil immersion objectives can potentially be used for OPM based 3D imaging in watery samples, owing to new insights into remote focusing [26].

Since the only moving parts are of low inertia (galvo mirrors), POPSIM has the potential for high-speed imaging that is camera or fluorescence limited. It could advantageously be combined with TIRF- or GI-SIM modalities. As such, clathrin mediated endocytosis could be monitored at the plasma membrane together with intracellular transport, just to name one application of such a hybrid modality.

We anticipate that POPSIM adds new spatiotemporal imaging capabilities to fluorescence microscopy, which may prove useful to follow rapid cellular dynamics, and may spur further developments in OPM, SIM and projection imaging.

## Acknowledgments

The authors are grateful to Jens C. Schmidt for providing the ER labeled cells and Madeleine Marlar-Pavey for preparing the mitochondria labeled cells. We are also grateful to Felix Zhou for help with the registration code.

## References

1. M. G. Gustafsson, “Surpassing the lateral resolution limit by a factor of two using structured illumination microscopy,” Journal of microscopy 198, 82–87 (2000).

2. M. G. Gustafsson, L. Shao, P. M. Carlton, C. R. Wang, I. N. Golubovskaya, W. Z. Cande, D. A. Agard, and J. W. Sedat, “Three-dimensional resolution doubling in wide-field fluorescence microscopy by structured illumination,” Biophysical journal 94, 4957–4970 (2008).

3. P. Kner, B. B. Chhun, E. R. Griffis, L. Winoto, and M. G. Gustafsson, “Super-resolution video microscopy of live cells by structured illumination,” Nature methods 6, 339–342 (2009).

4. Y. Guo, D. Li, S. Zhang, Y. Yang, J.-J. Liu, X. Wang, C. Liu, D. E. Milkie, R. P. Moore, and U. S. Tulu, “Visualizing intracellular organelle and cytoskeletal interactions at nanoscale resolution on millisecond timescales,” Cell 175, 1430–1442. e1417 (2018).

5. H.-W. Lu-Walther, M. Kielhorn, R. Förster, A. Jost, K. Wicker, and R. Heintzmann, “fastSIM: a practical implementation of fast structured illumination microscopy,” Methods and Applications in Fluorescence 3, 014001 (2015).

6. A. Sandmeyer, M. Lachetta, H. Sandmeyer, W. HuLJbner, T. Huser, and M. MuLJller, “Cost-effective live cell structured illumination microscopy with video-rate imaging,” ACS Photonics 8, 1639–1648 (2021).

7. L. Shao, P. Kner, E. H. Rego, and M. G. Gustafsson, “Super-resolution 3D microscopy of live whole cells using structured illumination,” Nature methods 8, 1044–1046 (2011).

8. R. Fiolka, L. Shao, E. H. Rego, M. W. Davidson, and M. G. Gustafsson, “Time-lapse two-color 3D imaging of live cells with doubled resolution using structured illumination,” Proceedings of the National Academy of Sciences 109, 5311–5315 (2012).

9. K. Wicker, O. Mandula, G. Best, R. Fiolka, and R. Heintzmann, “Phase optimisation for structured illumination microscopy,” Optics express 21, 2032–2049 (2013).

10. Y. Mo, K. Wang, L. Li, S. Xing, S. Ye, J. Wen, X. Duan, Z. Luo, W. Gou, and T. Chen, “Quantitative structured illumination microscopy via a physical model-based background filtering algorithm reveals actin dynamics,” Nature Communications 14, 3089 (2023).

11. A. Markwirth, M. Lachetta, V. Mönkemöller, R. Heintzmann, W. Hübner, T. Huser, and M. Müller, “Video-rate multi-color structured illumination microscopy with simultaneous real-time reconstruction,” Nature communications 10, 4315 (2019).

12. B. Chen, B.-J. Chang, P. Roudot, F. Zhou, E. Sapoznik, M. Marlar-Pavey, J. B. Hayes, P. T. Brown, C.- W. Zeng, T. Lambert, J. R. Friedman, C.-L. Zhang, D. T. Burnette, D. P. Shepherd, K. M. Dean, and R. P. Fiolka, “Resolution doubling in light-sheet microscopy via oblique plane structured illumination,” Nature Methods 19, 1419–1426 (2022).

13. B.-J. Chang, J. D. Manton, E. Sapoznik, T. Pohlkamp, T. S. Terrones, E. S. Welf, V. S. Murali, P. Roudot, K. Hake, L. Whitehead, A. G. York, K. M. Dean, and R. Fiolka, “Real-time multi-angle projection imaging of biological dynamics,” Nature Methods 18, 829–834 (2021).

14. S. Quirin, N. Vladimirov, C.-T. Yang, D. S. Peterka, R. Yuste, and M. B. Ahrens, “Calcium imaging of neural circuits with extended depth-of-field light-sheet microscopy,” Optics Letters 41(2016).

15. S. Abrahamsson, S. Usawa, and M. Gustafsson, “A new approach to extended focus for high-speed high-resolution biological microscopy,” in Three-Dimensional and Multidimensional Microscopy: Image Acquisition and Processing XIII, (SPIE, 2006), 128–135.

16. R. Tomer, M. Lovett-Barron, I. Kauvar, A. Andalman, Vanessa M. Burns, S. Sankaran, L. Grosenick, M. Broxton, S. Yang, and K. Deisseroth, “SPED Light Sheet Microscopy: Fast Mapping of Biological System Structure and Function,” Cell 163, 1796–1806 (2015).

17. W. J. Shain, N. A. Vickers, B. B. Goldberg, T. Bifano, and J. Mertz, “Extended depth-of-field microscopy with a high-speed deformable mirror,” Optics Letters 42, 995–998 (2017).

18. C. Dunsby, “Optically sectioned imaging by oblique plane microscopy,” Optics Express 16(2008).

19. P. Lacroute and M. Levoy, “Fast volume rendering using a shear-warp factorization of the viewing transformation,” in Proceedings of the 21st annual conference on Computer graphics and interactive techniques - SIGGRAPH ‘94, (1994), pp. 451–458.

20. P. T. Brown, R. Kruithoff, G. J. Seedorf, and D. P. Shepherd, “Multicolor structured illumination microscopy and quantitative control of polychromatic light with a digital micromirror device,” Biomedical Optics Express 12, 3700–3716 (2021).

21. A. Descloux, K. S. Grußmayer, and A. Radenovic, “Parameter-free image resolution estimation based on decorrelation analysis,” Nature methods 16, 918–924 (2019).

22. J. Nixon-Abell, C. J. Obara, A. V. Weigel, D. Li, W. R. Legant, C. S. Xu, H. A. Pasolli, K. Harvey, H. F. Hess, and E. Betzig, “Increased spatiotemporal resolution reveals highly dynamic dense tubular matrices in the peripheral ER,” Science 354, aaf3928 (2016).

23. L. K. Schroeder, A. E. Barentine, H. Merta, S. Schweighofer, Y. Zhang, D. Baddeley, J. Bewersdorf, and S. Bahmanyar, “Dynamic nanoscale morphology of the ER surveyed by STED microscopy,” Journal of Cell Biology 218, 83–96 (2019).

24. B.-C. Chen, W. R. Legant, K. Wang, L. Shao, D. E. Milkie, M. W. Davidson, C. Janetopoulos, X. S. Wu, J. A. Hammer, Z. Liu, B. P. English, Y. Mimori-Kiyosue, D. P. Romero, A. T. Ritter, J. Lippincott-Schwartz, L. Fritz-Laylin, R. D. Mullins, D. M. Mitchell, J. N. Bembenek, A.-C. Reymann, R. Böhme, S. W. Grill, J. T. Wang, G. Seydoux, U. S. Tulu, D. P. Kiehart, and E. Betzig, “Lattice light-sheet microscopy: Imaging molecules to embryos at high spatiotemporal resolution,” Science 346(2014).

25. B.-J. Chang, W.-C. Tang, Y.-T. Liu, Y.-C. Tsai, C. Tsao, P. Chen, and B.-C. Chen, “Two-beam interference lattice lightsheet for structured illumination microscopy,” Journal of Physics D: Applied Physics 53, 044005 (2019).

26. A. Millett-Sikking, “Any immersion remote refocus (AIRR) microscopy,” doi:10.5281/zenodo.7425649 (2022).

